# Impact of referencing scheme on decoding performance of LFP-based brain-machine interface

**DOI:** 10.1101/2020.05.03.075218

**Authors:** Nur Ahmadi, Timothy G. Constandinou, Christos-Savvas Bouganis

## Abstract

**Objective:** There has recently been an increasing interest in local field potential (LFP) for brain-machine interface (BMI) applications due to its desirable properties (signal stability and low bandwidth). LFP is typically recorded with respect to a single unipolar reference which is susceptible to common noise. Several referencing schemes have been proposed to eliminate the common noise, such as bipolar reference, current source density (CSD), and common average reference (CAR). However, to date, there have not been any studies to investigate the impact of these referencing schemes on decoding performance of LFP-based BMIs.

**Approach:** To address this issue, we comprehensively examined the impact of different referencing schemes and LFP features on the performance of hand kinematic decoding using a deep learning method. We used LFPs chronically recorded from the motor cortex area of a monkey while performing reaching tasks.

**Main results:** Experimental results revealed that local motor potential (LMP) emerged as the most informative feature regardless of the referencing schemes. Using LMP as the feature, CAR was found to yield consistently better decoding performance than other referencing schemes over long-term recording sessions.

**Significance:** Overall, our results suggest the potential use of LMP coupled with CAR for enhancing the decoding performance of LFP-based BMIs.

## 1. Introduction

Brain-machine interfaces (BMIs) have emerged as a promising technology to restore lost motor function in severely paralysed individuals due to neurological disorders such as amyotrophic lateral sclerosis (ALS), spinal cord injury (SCI), and stroke. BMIs allow these individuals to interact with the environment via assistive devices (e.g. robotic arm, computer cursor, functional electrical stimulator) controlled directly by their thoughts. In recent years, there has been an increasing interest in using local field potential (LFP) as an alternative or complementary input signal for BMIs [1, 2, 3]. This trend arises from the desirable properties of LFP: signal stability and low bandwidth. LFPs have been demonstrated to yield long-term decoding stability even in the absence of action potential (spike) activity [1, 4, 5]. Additionally, LFPs can be sampled and processed at much lower sampling rate than spike, which translates into lower power consumption, thereby minimising the risk of overheating, reducing the form factor and increasing lifespan of a BMI device [1, 3]. Many researchers are thus interested in attempting to extract informative features for decoding kinetic or kinematic parameters of movement [6, 7, 2, 8].

A great deal of literature has been published on the use of LFPs for BMIs in non-human primates [9, 10, 1, 4, 5, 2, 3] and humans [11, 12, 13, 14]. However, most LFP-based BMIs use neural signals recorded relative to a single common reference (hereinafter referred to as unipolar reference) which are susceptible to common noise due to different sources including non-silent reference and volume conduction [15, 16]. Since there is no reference site in the body free from electrical activity [17] and neural signals are measured as the potential differences between the active electrodes and reference, electrical activity in the reference will affect the recordings in all active electrodes. In addition, electrical activity from distant sources can also contaminate the recordings through the volume conduction mechanism. This common noise may degrade LFP’s signal-to-noise ratio (SNR) and lead to suboptimal BMI decoding performance. Several referencing schemes have been proposed to remove the common noise, such as bipolar reference, common source density (CSD), and common average reference (CAR). Previous studies have assessed the impact of these referencing schemes on the analyses of power, phase, coherence, and Granger causality in LFPs [15, 16, 18, 19, 20]. So far, however, there is no study that examines the impact of these referencing schemes on decoding performance of LFP-based BMIs.

This present study aims to address the above knowledge gap by systematically evaluating the impact of different referencing schemes on decoding performance of LFP-based BMIs. We used long-term LFP data that were chronically recorded from the motor cortex area of a macaque monkey (*Macaca mulatta*) while performing self-paced reaching tasks. We employed a deep learning method called long short-term memory (LSTM) network for decoding hand kinematics from LFPs. Firstly, We examined the informativeness of different LFP features across referencing schemes. Secondly, we assessed the impact of interelectrode distance in bipolar reference on decoding performance. We then compared the decoding performance across referencing schemes. Lastly, we investigated the impact of referencing schemes on interchannel correlation of LFPs.

The remainder of this paper is organised as follows. Section 2 describes the dataset, methodology, and experimental setup. Section 3 presents the empirical results, followed by discussion and analyses in Section 4. Finally, the conclusion is drawn in Section 5.

## 2. Methods

A schematic overview of LFP-based BMI system used in this study is illustrated in figure 1. All the experiments and analyses were done in Python. Keras/TensorFlow deep learning framework [21] was used to implement the decoding algorithms.

**Figure 1.**
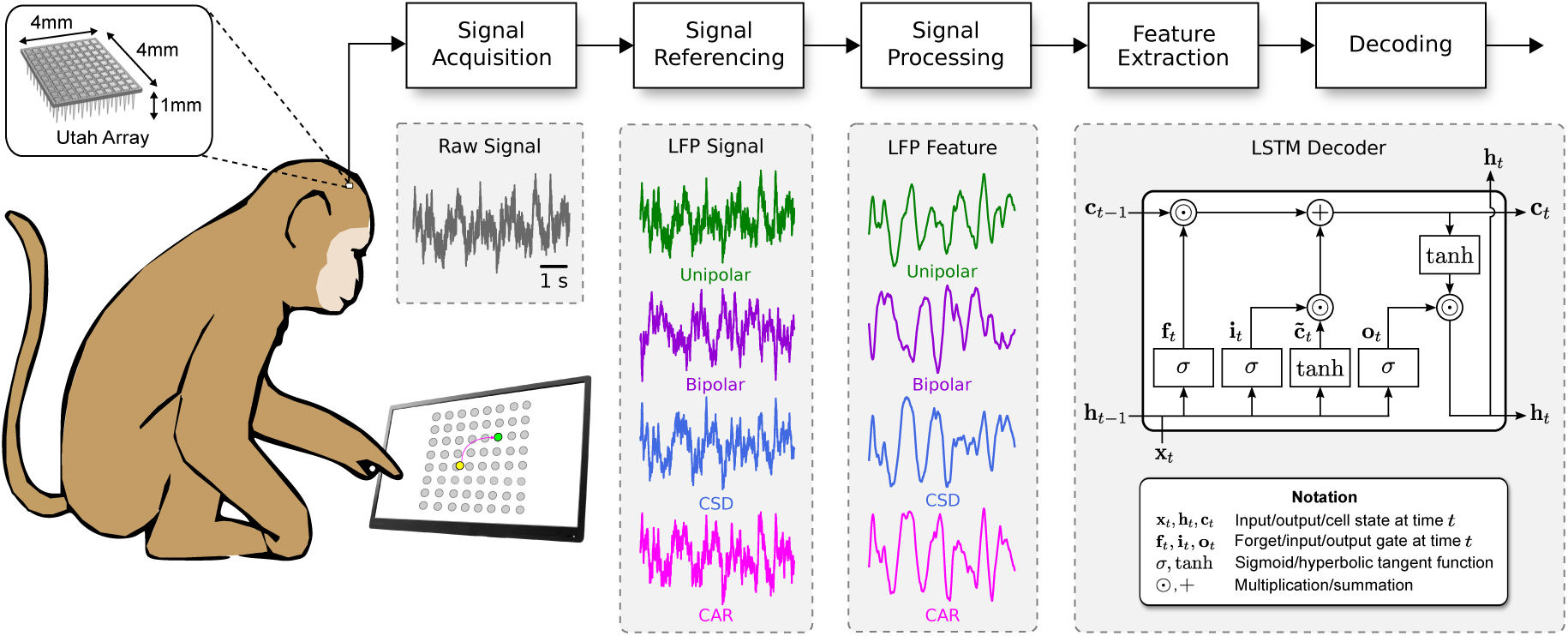
Schematic overview of LFP-based BMI system using LSTM decoder with different referencing schemes.

### 2.1. Signal acquisition

We used an existing, freely-available neural dataset deposited by Sabes lab [22]. This dataset contains long-term raw neural data and the corresponding behavioural task, allowing us to analyse the impact of the referencing schemes over a long period of time. A total of 26 recording sessions were included for experiments and analyses, spanning over 7.3 months between the first (I20160627_01) and last (I20170131_02) sessions, with an average duration of 8.88 ± 1.96 minutes. The neural signals were recorded from the primary motor cortex (M1) area of an adult male Rhesus macaque monkey (*Macaca mulatta*) with a 96-channel Utah microelectrode array (platinum coating, 1 mm electrode length, 400 *µ*m spacing, 400 kΩ impedance). The recordings were pre-filtered using a 4th-order low-pass filter with a roll-off of 24 dB per octave at 7.5 kHz and sampled in 16-bit resolution at 24.4 kHz. The digitised recordings are hereinafter called as raw neural signals. Details on the data acquisition setup can be found elsewhere [23]. Due to the fixed length of Utah array, the analysis in this study is confined to signals recorded at a specific depth. Therefore, referencing schemes are compared on 2D spatial properties parallel to the cortical surface.

### 2.2. Behavioural task

The neural recordings were acquired while the monkey was performing a point-to-point task, that is, to reach randomly drawn circular targets uniformly distributed around an 8-by-8 square grid. A sequence of new random targets was presented immediately and continuously after target acquisition without an inter-trial interval. Fingertip position in *x*-*y* coordinates was sampled at 250 Hz and non-causally filtered with a 4th order Butterworth low-pass filter at 10 Hz to reject sensor noise. Fingertip velocity was then computed from the position data by using a discrete derivative. A more detailed description of the behavioural task setup is given in [23].

### 2.3. Signal referencing

We used four referencing schemes namely unipolar reference, bipolar reference, current source density (CSD), and common average reference (CAR), each of which is described as follows.

#### 2.3.1. Unipolar reference

Unipolar reference in the form of a silver wire placed under the dura (several cm away from the electrodes) was originally used during neural signal acquisition. The recorded neural signal at electrode *i*, represented by 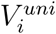, is measured as:

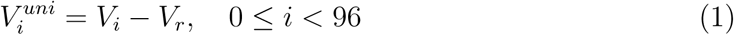

where *V*_*i*_ and *V*_*r*_ denote the electric potential at the electrode *i* and reference, respectively. In total there was 96 unipolar signals (i.e. channels). The characteristics of the recording setup, e.g. very low added noise by instrumentation, high-resolution data conversion, and broadband data, allow us to emulate other referencing scheme in software. All filtering operation was performed digitally in forward and backward directions to avoid any phase distortion.

#### 2.3.2. Bipolar reference

Bipolar referencing, initially used in EEG studies [24], was obtained by computing the difference between a pair of unipolar signals as follows:

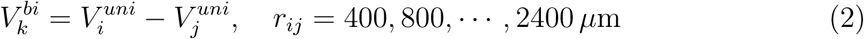

where *r*_*ij*_ denotes a predefined distance between electrode *i* and electrode *j* ranging from 400 *µ*m to 2400 *µ*m with an increment of 400 *µ*m, which yielded a total of 172, 152, 132, 112, 172, and 72 bipolar signals, respectively.

#### 2.3.3. Current source density (CSD)

CSD, also known as Laplacian referencing [25], was computed by subtracting the average of four nearest neighbouring electrodes with equal distance of 400 *µ*m as formulated below:

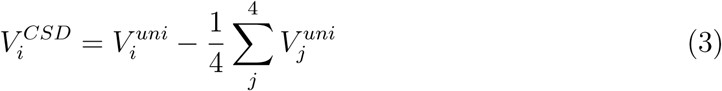

Only electrodes having 4 equidistant neighbours were included for further experiments and analyses, yielding a total of 64 CSD signals (i.e. electrodes at the edge were excluded).

#### 2.3.4. Common average reference (CAR)

Originally used for ECG [26] in 1930s and firstly adopted for EEG in 1950s [27], CAR has become one of the most popular referencing schemes. CAR was obtained by subtracting the average of all the unipolar signals as described by the following equation:

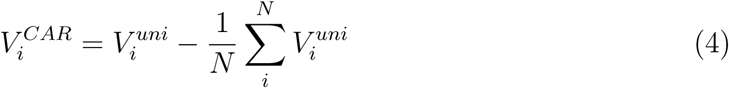

CAR referencing resulted in the same number of signals as the original unipolar reference (96 channels).

### 2.4. Signal processing

To obtain LFP signals, the raw neural signals were low-pass filtered with a 4th-order Butterworth filter at 300 Hz and then downsampled to 1 kHz. The filtering was performed on the forward and backward directions to avoid any phase shift. All the LFP channels were included for experiments and analyses.

### 2.5. Feature extraction

For each LFP channel, six different features were extracted (one feature in the time domain and five features in the frequency domain). The time-domain feature was an average amplitude called local motor potential (LMP) whereas the frequency domain features were average spectral power in five different frequency bands: delta (1–4 Hz), theta (3–10 Hz), alpha (12–23 Hz), beta (27–38 Hz) and gamma (50–300 Hz). The selection of these frequency bands followed a previous study by Stavisky *et al.* [2]. The LMP was obtained by using a moving average filter with 256 ms rectangular window. To extract the spectral power features, we first band-pass filtered the LFP signals using 3rd-order Butterworth filter at each frequency band and then applied short-time Fourier transform (STFT) with a 256 ms Hanning window. All the feature extractions were performed in an overlapping fashion (252 ms overlap) to yield a sample every 4 ms (matching the sampling rate of the kinematic data).

### 2.6. Decoding

#### 2.6.1. Long short-term memory (LSTM) decoder

LSTM, proposed by Hochreiter and Schmidhuber in 1997 [28], has become one of the most popular and widely used deep learning methods. LSTMs have demonstrated state-of-the-art performances in various applications such as language modelling and translation, speech recognition and synthesis, and video classification and captioning [29]. Our previous study has also shown that LSTM can achieve higher decoding performance when compared to a Kalman filter [30]. LSTM successfully addresses the vanishing gradient problem commonly encountered in traditional recurrent neural networks (RNNs) and is capable of learning long-term temporal dependencies. LSTM maintains its internal states overtime and controls the flow of information via a memory cell and gating mechanism, as illustrated in figure 1. The LSTM states at each timestep *t* are described by the following equations:

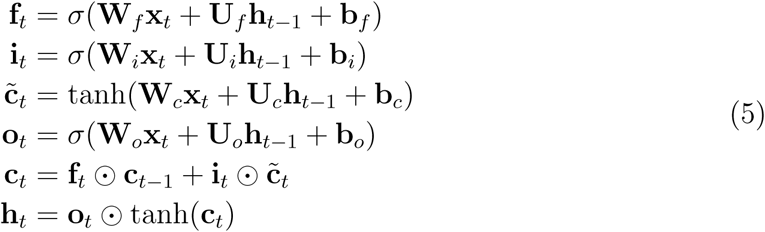

where **x, h, f**, **i, o, c** denote the input, output, forget gate, input gate, output gate, and memory cell, respectively. *σ*, tanh, and ⊙ represent consecutively logistic sigmoid, hyperbolic tangent, and element-wise multiplication. Weight matrices (**W, U**) and bias vectors (**b**) are the learnable parameters.

We used one layer LSTM and set empirically the timestep (i.e. sequence length) hyperparameter to 2. As described in the next subsection, hyperparameter optimisation technique was used to optimise other hyperparameters which include the number of units, number of epochs, batch size, dropout rate and learning rate. To obtain the final output, the last timestep of the LSTM states were connected to a fully connected layer with a linear activation function.

#### 2.6.2. LSTM decoder training and optimisation

The dataset was firstly divided into 10 non-overlapping contiguous blocks of equal size which were categorised further into three sets: training (8 concatenated blocks), validation (1 block) and testing (1 block). The training set was used to train the model; the validation set was used to find the optimised hyperparameters; the testing set was used to evaluate the optimised model. Next, the LFP features were standardised (i.e. z-transformed) to zero mean and unit variance, whereas the kinematic data were normalised to zero mean. The model was trained using RMSprop optimiser and root mean squared error (RMSE) loss function. Bayesian optimisation package called Hyperopt [31] was used to find the best performing hyperparameters from predefined value ranges. The resulting optimised hyperparameters for each LFP feature and referencing scheme are given in Table 1 and Table 2, respectively. As hyperparameter optimisation took considerably long computation time, it was performed only in the first recording session; for the subsequent sessions, we used the same hyperparameter configuration.

**Table 1.** Hyperparameter configuration of LSTM decoders across LFP features

| Hyperparameter | Value Range | LFP Feature |  |  |  |  |  |
| --- | --- | --- | --- | --- | --- | --- | --- |
|  |  | LMP | Delta | Theta | Alpha | Beta | Gamma |
| Number of units | $\{50, 75, \dots, 200\}$ | 175 | 175 | 100 | 75 | 175 | 200 |
| Number of epochs | $\{2, 3, \dots, 8\}$ | 3 | 2 | 2 | 2 | 2 | 2 |
| Batch size | $\{64, 96, 128\}$ | 64 | 64 | 64 | 64 | 64 | 64 |
| Dropout rate | $\{0, 0.1, \dots, 0.5\}$ | 0.4 | 0.2 | 0.2 | 0.3 | 0.2 | 0.3 |
| Learning rate | $\{5, 10, \dots, 50\} \times 10^{-4}$ | 0.0005 | 0.001 | 0.0005 | 0.0005 | 0.0005 | 0.0005 |

**Table 2.** Hyperparameter configuration of LSTM decoders across referencing schemes.

| Hyperparameter | Value Range | Referencing scheme |  |  |  |
| --- | --- | --- | --- | --- | --- |
|  |  | Unipolar | Bipolar | CSD | CAR |
| Number of units | $\{50, 75, \dots, 200\}$ | 175 | 150 | 50 | 175 |
| Number of epochs | $\{2, 3, \dots, 8\}$ | 3 | 3 | 3 | 5 |
| Batch size | $\{64, 96, 128\}$ | 64 | 96 | 96 | 64 |
| Dropout rate | $\{0, 0.1, \dots, 0.5\}$ | 0.4 | 0.5 | 0.4 | 0.4 |
| Learning rate | $\{5, 10, \dots, 50\} \times 10^{-4}$ | 0.0005 | 0.0045 | 0.001 | 0.0045 |

### 2.7. Performance evaluation and metrics

Decoding performance of each model was evaluated using two commonly used metrics: root mean square error (RMSE) and Pearson’s correlation coefficients (CC) [9, 10, 32]. RMSE measures the average magnitude of the decoding error, whereas CC measures the linear correlation between the true and decoded hand kinematics. RMSE and CC are computed using the following formula:

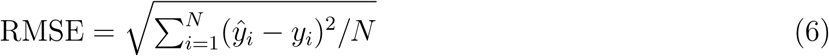

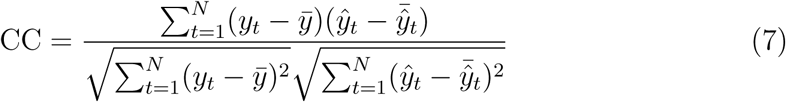

where *y*_*t*_ and 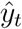 represent the true and decoded hand kinematics at timestep *t*, respectively and *N* denotes the total number of samples.

We calculated the mean and confidence interval of the decoding performance for each session (*n* = 10 blocks) and across all sessions (*n* = 26 sessions). Statistical significant tests between a pair of different LFP features or different referencing schemes were conducted using a paired two-tailed *t*-test whenever the difference between the pairs follows a normal distribution, otherwise using a Wilcoxon signed-rank test. The significance level (*α*) was 0.05.

## 3. Results

### 3.1. LFP feature comparison and selection

Firstly, we assessed the informativeness of LFP features by measuring their decoding performance under unipolar reference using independently optimised LSTM decoders. Experimental results revealed that LMP yielded the highest decoding performance (smallest RMSE and largest CC), followed by gamma, delta, theta, alpha, and beta, respectively (see figure 2). The circle mark and horizontal line within each boxplot denote the mean and median, respectively. The solid coloured box represents the interquartile range between 25th and 75th percentiles. The whisker extends to 1.5 times the interquartile range. As shown in figures 2(a) and 2(c), LMP and gamma were the two most informative features, yielding RMSE = 45.65 ± 0.90 and CC = 0.81 ± 0.01 for LMP and RMSE = 60.62 ± 1.50 and CC = 0.61 ± 0.01 for gamma. In this study, RMSE and CC performance are written in terms of mean ± standard error of the mean (SEM) unless otherwise noted. Relative to gamma, the performance of LMP in terms of RMSE and CC improved on average by 24.6% and 32.6%, respectively, as can be seen from figures 2(b) and 2(d). The performance of other LFP features (delta, theta, alpha, and beta) degraded on average by 15.8%–26.9% (RMSE) and 29.7%–89.1% depending on the feature.

**Figure 2.**
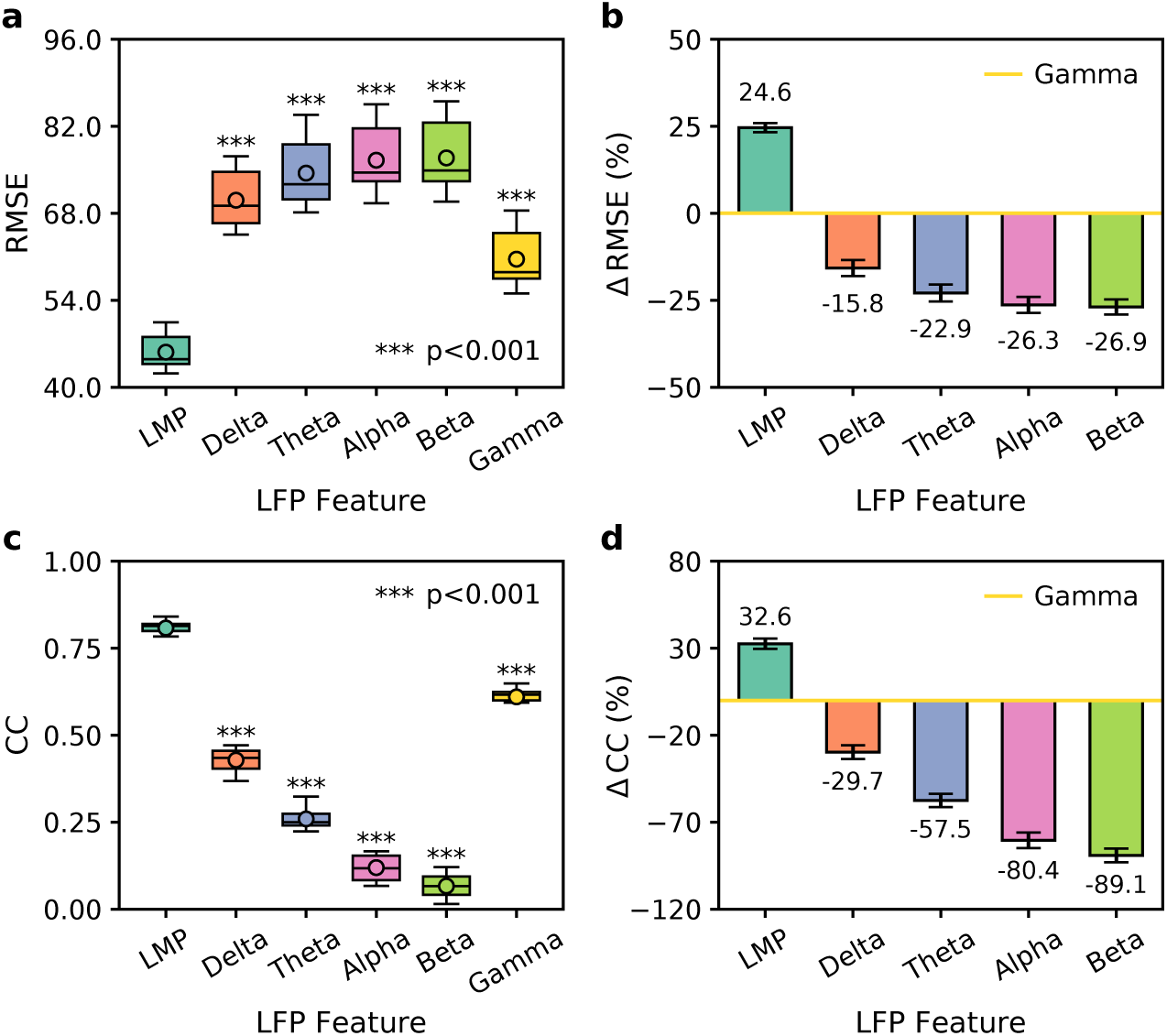
Decoding performance comparison across different LFP features in unipolar reference (data from recording session I20160627_01). (a), (c) Boxplot comparison across features measured in RMSE and CC, respectively. Asterisks indicate features that had statistically significant different performances from that of LMP (*** p < 0.001). (b), (d) Performance improvement/degradation (in percent RMSE and CC, respectively) relative to gamma. Positive (negative) value indicates performance improvement (degradation). Black error bars denote 95% confidence intervals.

We then examined whether the same trend was also observed from different referencing schemes. Results from bipolar reference showed the same order of feature informativeness as of unipolar reference (in descending order): LMP > gamma > delta > theta > alpha > beta. There were some variations in the order of feature informativeness in the cases of CSD and CAR, which were LMP > gamma > theta > delta > beta > alpha and LMP > gamma > delta > theta > beta> alpha, respectively. These extensive results revealed that LMP and gamma consistently emerged as the first and second most informative features. Therefore, for subsequent experiments, we selected to use only these two features.

### 3.2. Impact of interelectrode distance in bipolar reference

Next, we investigated the impact of interelectrode distance in bipolar reference by comparing the decoding performance from varying distances (400, 800, …, 2400 *µ*m) over long-term recording sessions. Figures 3 and 4 display the decoding performance comparison across different interelectrode distances over 26 sessions from LMP and gamma features, respectively. In the case of LMP, the decoding performance from the distances of 400, 800, 1200, and 2000 *µ*m (with exceptions from the distances of 1600 and 2400 *µ*m) were similar as can be observed in figures 3(a) and 3(b). The average RMSE and CC for the distance of 400 *µ*m were 49.64 ± 0.88 and 0.76 ± 0.01, respectively (figures 3(c) and 3(e)). Increasing the distances from 400 *µ*m to 2400 *µ*m did not improve the decoding performance, but, on the contrary, it tended to degrade the decoding performance as clearly visible in figures 3(d) and 3(f). Specifically, the decoding performance from the distances of 1600 *µ*m and 2400 *µ*m were statistically significantly lower than that of 400 *µ*m (*** p < 0.001). With respect to 400 *µ*m, the performance of 1600 *µ*m degraded by 4.6% (RMSE) and 3.7% (CC), whereas the performance of 2400 *µ*m degraded by 10.2% (RMSE) and 8.5% (CC).

**Figure 3.**
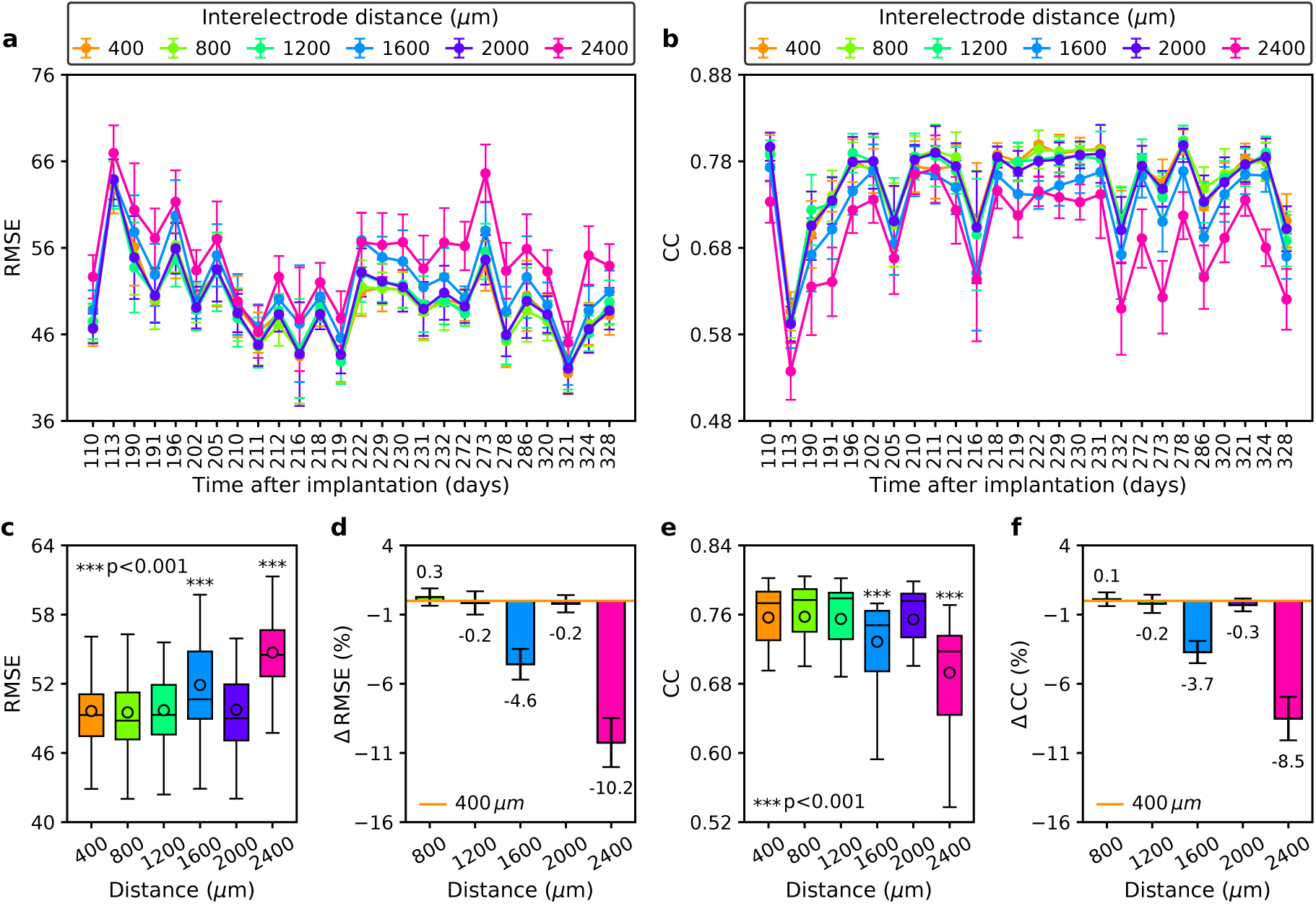
Decoding performance comparison across different interelectrode distances in bipolar reference with LMP feature. (a), (b) Performance comparison over long-term recording sessions measured in RMSE and CC, respectively. (c), (e) Boxplot comparison across distances and sessions measured in RMSE and CC, respectively. Asterisks indicate distances whose performances were statistically significant difference from that of 400 *µ*m (*** p < 0.001). (d), (f) Performance improvement/degradation (in percent RMSE and CC, respectively) relative to that of 400 *µ*m. Positive (negative) value indicates performance improvement (degradation). Black error bars denote 95% confidence intervals.

**Figure 4.**
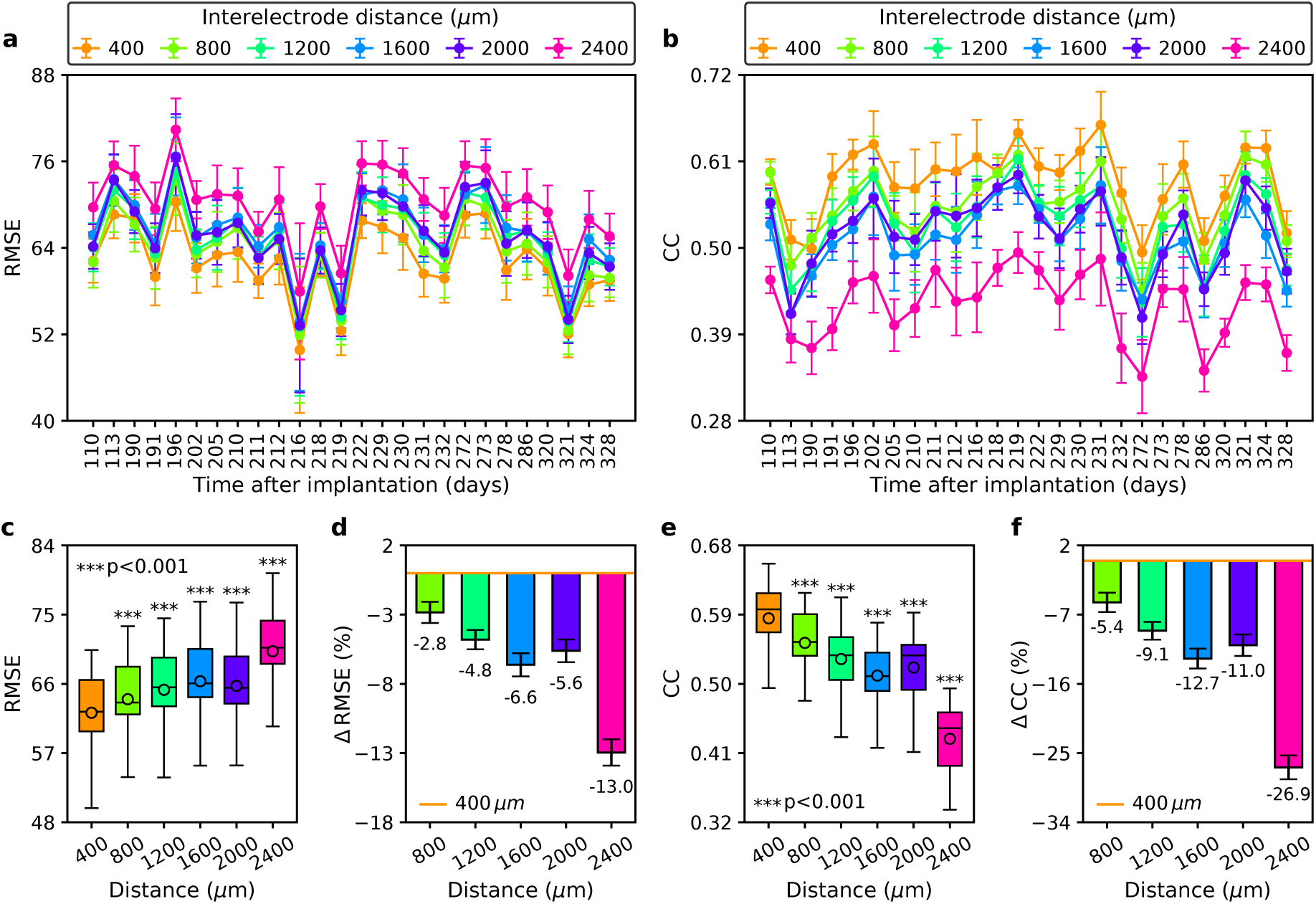
Decoding performance comparison across different interelectrode distance in bipolar reference with gamma feature. (a), (b) Performance comparison over long-term recording sessions measured in RMSE and CC, respectively. (c), (e) Boxplot comparison across distances and sessions measured in RMSE and CC, respectively. Asterisks indicate distances whose performances were statistically significant difference from that of 400 *µ*m (*** p < 0.001). (d), (f) Performance improvement/degradation (in percent RMSE and CC, respectively) relative to that of 400 *µ*m. Positive (negative) value indicates performance improvement (degradation). Black error bars denote 95% confidence intervals.

In the case of gamma, the average decoding performance of 400 *µ*m was 62.24 ±1.03 (RMSE) and 0.59 ± 0.01 (CC) and was consistently better than that of larger distances across sessions as illustrated in figures 4(a) and 4(b). Statistical significant tests showed that the decoding performance of larger distances exhibited significant difference than that of 400 *µ*m (*** p < 0.001) as evident from figure 4(c) for RMSE and figure 4(e) for CC. Although in essence similar to the case of LMP, the impact of increasing interelectrode distances was more clearly observed in the case of gamma. As shown in figures 4(d) and 4(f), increasing the distances resulted in performance degradation of 2.8%–13.0% (RMSE) and 5.4%–26.9% (CC) relative to that of 400 *µ*m. We thus selected an interelectrode distance of 400 *µ*m for subsequent experiments.

### 3.3. Impact of referencing scheme on decoding performance

We then evaluated the impact of different referencing schemes on decoding performance of LSTM decoder driven by LMP feature. The decoding performance comparison across referencing schemes from LMP-driven LSTM decoder over 26 recording sessions measured in RMSE and CC are illustrated in figures 5(a) and 5(b), respectively. The decoding performance of each referencing scheme averaged across sessions was as follows: unipolar (RMSE = 50.97 ± 0.96, CC = 0.76 ± 0.01), bipolar (RMSE = 49.64 ± 0.88, CC = 0.76 ± 0.01), CSD (RMSE = 61.72 ± 1.11, CC = 0.61 ± 0.01), and CAR (RMSE = 48.08 ± 0.97, CC = 0.77 ± 0.01). Sorting the referencing schemes descendingly according to their decoding performance resulted in the following order: CAR > bipolar > unipolar > CSD (see figures 5(c) and 5(e)). Statistical tests showed that, compared to that of unipolar, the performance of CSD and CAR had significant differences in both RMSE and CC metrics (*** p < 0.001). CSD resulted in average performance degradation of 21.2% (RMSE) and 19.1% (CC), while CAR yielded an average performance improvement of 5.6% (RMSE) and 1.9% (CC) as can be seen in figures 5(d) and 5(f). The performance of bipolar, however, only showed a significant difference in RMSE metric (** p < 0.01). Snippet examples of decoded *x*- and *y*- velocities along with their ground truth values from four referencing schemes are illustrated in figure 5(g)-(j).

**Figure 5.**
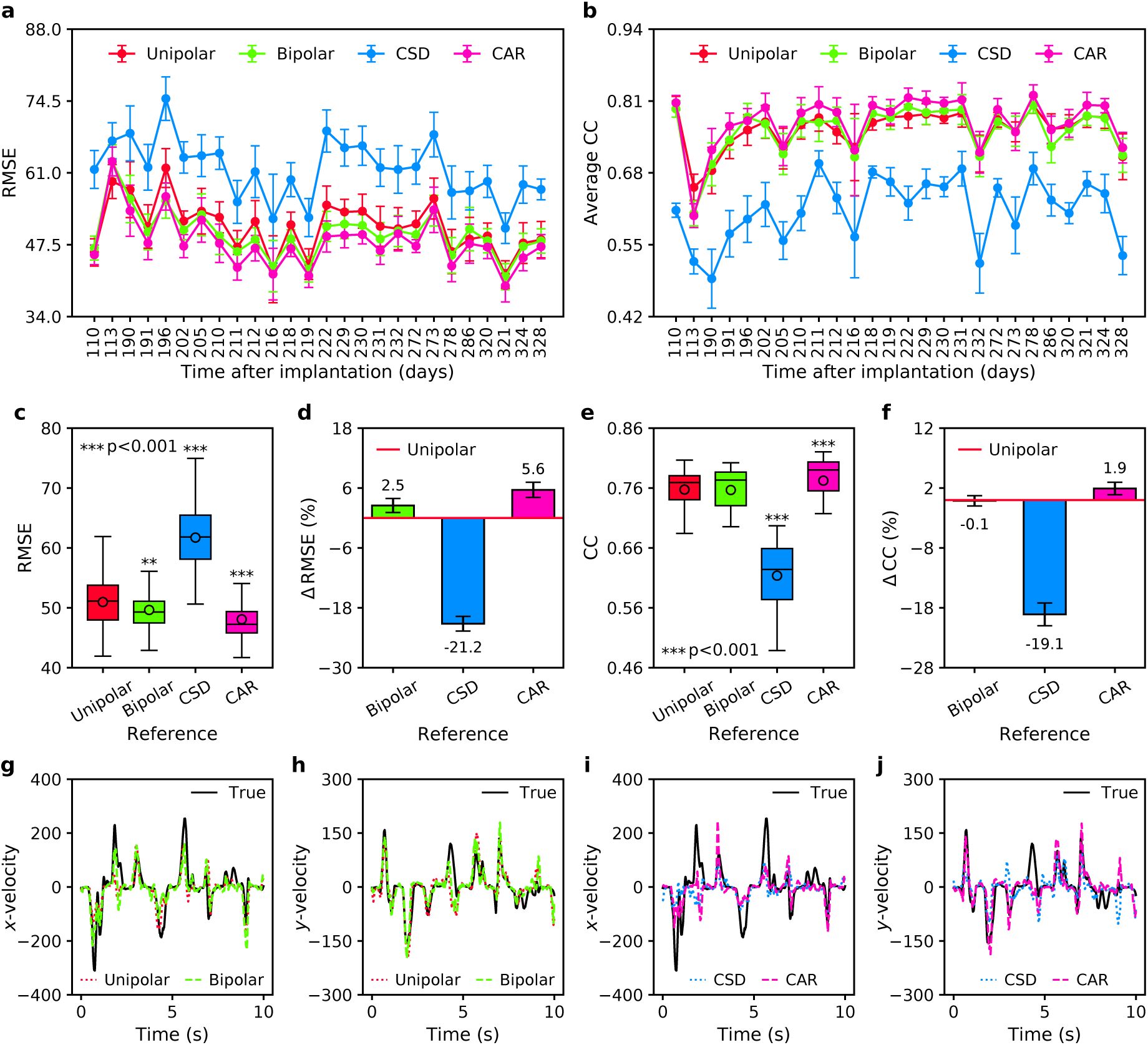
Decoding performance comparison across different referencing schemes using LMP-driven LSTM decoder. (a), (b) Performance comparison over long-term recording sessions measured in RMSE and CC, respectively. (c), (e) Boxplot comparison across sessions measured in RMSE and CC, respectively. Asterisks indicate referencing schemes that yielded statistically significant different performances from that of unipolar reference (** p < 0.01, *** p < 0.001). (d), (f) Performance improvement/degradation (in percent RMSE and CC, respectively) relative to unipolar reference. Positive (negative) value indicates performance improvement (degradation). Black error bars denote 95% confidence intervals. (g)-(j) Snippet examples of true and decoded velocities in *x*- and *y*- coordinates from different referencing schemes (data from recording session I20160627_01).

To check whether a similar trend was observed when using gamma feature, we compared the decoding performance of gamma-driven LSTM decoder across referencing schemes. The result of RMSE and CC performance comparison over 26 recording sessions were depicted in figures 6(a) and 6(b), respectively. In comparison to the case of LMP, here we found contrasting results when using bipolar and CAR, but consistent result when using CSD. The order of decoding performance across referencing schemes (from high to low) was bipolar (RMSE = 62.24 ± 1.03, CC = 0.59 ± 0.01), CAR (RMSE = 64.03 ± 1.12, CC = 0.55 ± 0.01), unipolar (RMSE = 63.88 ± 1.10, CC = 0.55 ± 0.01), and CSD (RMSE = 68.11 ± 1.11, CC = 0.46 ± 0.01). With respect to unipolar, bipolar and CSD yielded significant differences in the decoding performance (*** p < 0.001), whereas CAR did not differ significantly as can be seen in figures 6(c) and 6(e). Relative to unipolar, bipolar improved the average decoding performance by 2.5% (RMSE) and 6.7% (CC), while CSD degraded the average decoding performance by 6.7% (RMSE) and 16.9% (CC) as plotted in figures 6(d) and 6(f). It is worth noting that the decoding performance from gamma was consistently lower than that from LMP regardless of the referencing scheme.

**Figure 6.**
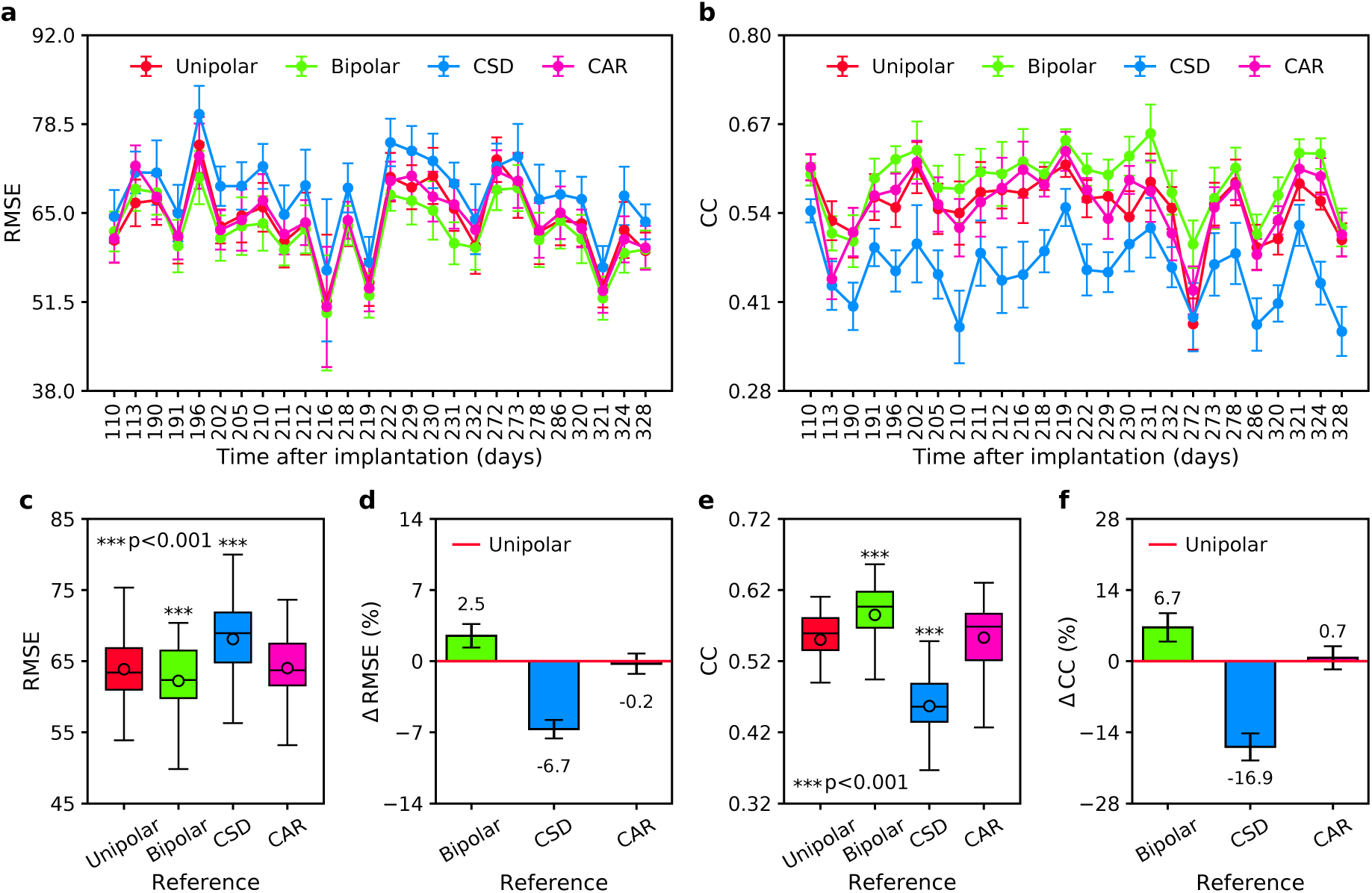
Decoding performance comparison across different referencing schemes using gamma-driven LSTM decoder. (a), (b) Performance comparison over long-term recording sessions measured in RMSE and CC, respectively. (c), (e) Boxplot comparison across sessions measured in RMSE and CC, respectively. Asterisks indicate referencing schemes that yielded statistically significant different performances from that of unipolar reference (*** p < 0.001). (d), (f) Performance improvement/degradation (in percent RMSE and CC, respectively) relative to unipolar reference. Positive (negative) value indicates performance improvement (degradation). Black error bars denote 95% confidence intervals.

We next sought to assess whether the impact of the referencing schemes on decoding performance remain the same when using different decoding algorithm. We thus employed a new decoder based on gated recurrent unit (GRU) network. GRU, proposed by Cho *et al.* in 2014 [33], uses an update gate and a reset gate to control information that flows into and out of a memory cell. GRU is similar to LSTM, but it has fewer parameters due to the absence of output gate. We compared the decoding performance across referencing schemes using LMP-driven GRU decoder. Figures 7(a) and 7(b) illustrate the decoding performance comparison over 26 sessions in terms of RMSE and CC, respectively. We observed the same pattern of decoding performance across referencing schemes as in the case of LMP-driven LSTM decoder. Sorted from high to low decoding performance, we obtained the following order: CAR (RMSE = 48.62±0.96, CC = 0.77±0.01), bipolar (RMSE = 50.00±0.89, CC = 0.75±0.01), unipolar (RMSE = 52.19 ± 0.86, CC = 0.73 ± 0.01), and CSD (RMSE = 61.86 ± 1.06, CC = 0.58 ± 0.01) as depicted in figures 7(c) and 7(e). Relative to unipolar, bipolar yielded performance improvement of 4.2% (RMSE) and 3.5% (CC), whereas CAR yielded performance improvement of 6.9% (RMSE) and 5.6% (CC) as shown in figures 7(d) and 7(f). On the other hand, the performance of CSD degraded by 18.6% (RMSE) and 20.3% (CC). Although the same pattern was observed between LMP-driven GRU and LMP-driven LSTM decoders, the magnitude of the decoding performance (RMSE and CC) obtained from LSTM were consistently better than that of GRU across all referencing schemes.

**Figure 7.**
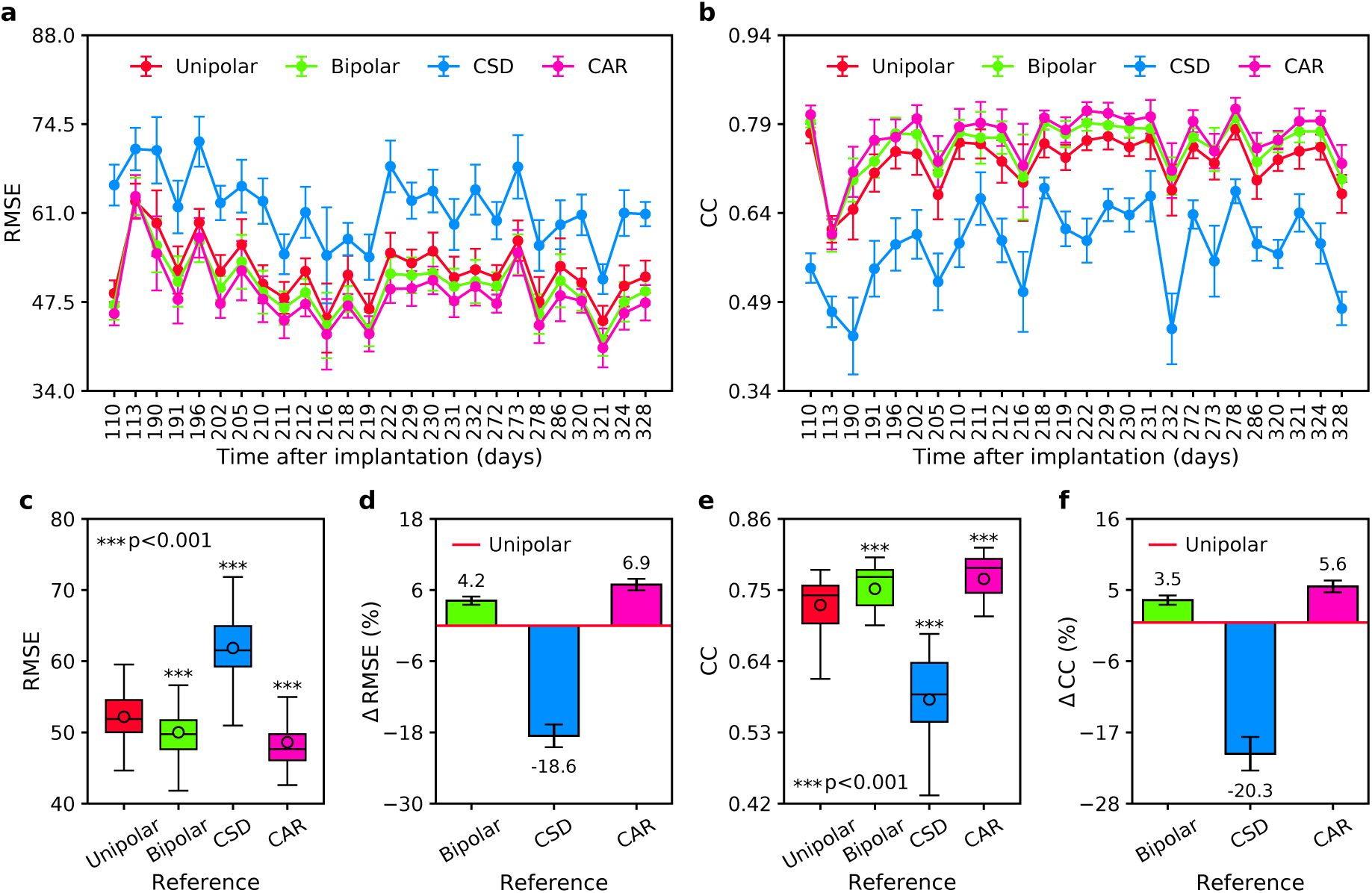
Decoding performance comparison across different referencing schemes using LMP-driven GRU decoder. (a), (b) Performance comparison over long-term recording sessions measured in RMSE and CC, respectively. (c), (e) Boxplot comparison across sessions measured in RMSE and CC, respectively. Asterisks indicate referencing schemes that yielded statistically significant different performances from that of unipolar reference (*** p < 0.001). (d), (f) Performance improvement/degradation (in percent RMSE and CC, respectively) relative to unipolar reference. Positive (negative) value indicates performance improvement (degradation). Black error bars denote 95% confidence intervals.

### 3.4. Impact of referencing scheme on interchannel correlation

Lastly, we examined the impact of referencing scheme on the interchannel correlation of LFPs by computing Pearson’s correlation coefficient among all unique possible pair combinations of LFP channels. Comparison of interchannel correlation across referencing schemes over 26 recording sessions is plotted in figure 8(a). Our results showed that, in comparison to unipolar, other referencing schemes yielded significantly lower average interchannel correlation (*** p < 0.001). By sorting the average interchannel correlation descendingly, we obtained the following order: unipolar (0.56±0.10) > CAR (0.20±0.07) > CSD (0.12±0.04) > bipolar (0.09±0.02). Figure 8(b) illustrates heatmaps of interchannel correlation matrix from each of referencing schemes.

**Figure 8.**
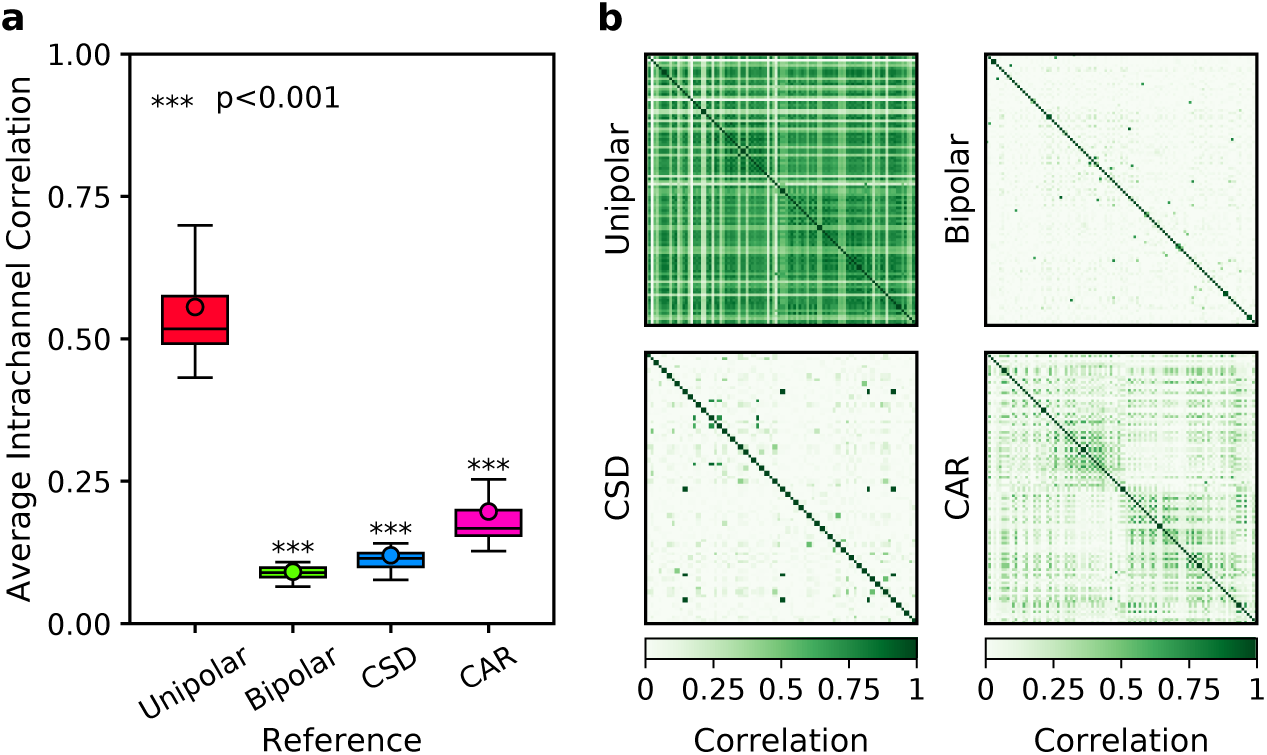
Comparison of interchannel correlation of LFPs across different referencing schemes. (a) Boxplot comparison of average interchannel correlation over 26 recording sessions. Asterisks indicate referencing schemes that yielded statistically significant different performances from that of unipolar reference (*** p < 0.001). (b) Comparison of interchannel correlation matrix across referencing schemes (data from recording session I20160627_01).

## 4. Discussion

We conducted an experimental investigation of the impact of four referencing schemes, namely unipolar, bipolar, current source density (CSD), and common average reference (CAR), on the decoding performance of LFP-based BMI. Firstly, we sought to determine which feature within LFP carries the highest movement-related information as characterised by its decoding performance. Several prior studies showed that power in the high-frequency band (i.e. high-gamma) of LFPs carried the highest information about movement compared to other frequency bands [9, 32, 34]. The high-gamma power is believed to reflect synchronised spiking activity arising from a local population of neurons near the recording electrodes [35, 36]. In these studies, however, local motor potential (LMP), which is a time-domain LFP feature, was not included in the experiments and analyses.

Our present study extends the above studies by comparing six LFP features consisting of LMP and power within five frequency bands: delta, theta, alpha, beta, and gamma. Our empirical results revealed that LMP significantly outperformed all the other LFP features. It was observed that LMP consistently achieved higher decoding accuracy across different referencing schemes. The high informativeness of LMP could be related to the view that LMP represents evoked LFP changes in response to behavioural tasks [37, 36]. Hitherto, it remains unclear what the physiological origin of LMP is. Different hypotheses have suggested that LMP is associated with firing rate modulation of neuronal population in the vicinity of the recording electrode [37] or activity of a distant neuronal population connected to the recording site [38, 39].

We also found that gamma always yielded the second-highest decoding performance after LMP. These findings are in good agreement with Stavisky *et al.*’s study [2], but contradict Flint *et al.*’s study [1] which showed that gamma was more informative than LMP. This contradiction could be due to the difference in the way we divide the LFP frequency bands and measure the informativeness of LFP features. Following Stavisky *et al.*’s study [2], we define gamma frequency band as 50–300 Hz, while Flint *et al.* split gamma frequency band into 70–200 Hz (low-gamma) and 200–300 Hz (high-gamma) [1]. In our study and [2], the informativeness of LFP feature is measured independently according to its decoding performance, whereas in [1] it is calculated based on relative contribution (i.e. filter weight) from top 150 mixed LFP features. Regardless of this contradiction, these three studies support the notion that LMP and gamma are both highly predictive for decoding kinematic data.

Next, we examined the impact of different interelectrode distances (400, 800, …, 2400 *µ*m) on the decoding performance in bipolar reference. Results from both LMP and gamma features showed that increasing the interelectrode distance tended to degrade the decoding performance. One possible reason for this trend could be related to the spatial spread of LFP which has been reported within around 200–400 *µ*m [40, 41, 42], although other studies suggest a larger spatial spread up to a few millimetres [43, 44]. Bipolar referencing is computed by subtracting one electrode from the other in an attempt to remove common noise component, yielding a more localised neural activity. Common noise arising from a distant source through volume conduction can be effectively removed if the interelectrode distance is sufficiently close [15]. If this is too close (high correlation between a pair of electrodes), bipolar referencing may pose a risk of subtracting out signal from approximately the same source. However, if this is too far apart, bipolar referencing could subtract out signal from a different source with a very little common noise component. An interelectrode distance of 400 *µ*m may reflect sufficiently different signal sources between the electrode pairs but exhibit considerably high common noise. This common noise could be effectively eliminated through bipolar referencing, which in turn lead to better decoding performance.

We then compared the impact of referencing schemes on the decoding performance using LMP-driven LSTM decoder. Results showed that CAR consistently outperformed all the other referencing schemes over long-term recording sessions. There were statistically significant differences in the decoding performance between CAR and other referencing schemes. The high decoding performance of CAR may be attributed to the fact that we use a relatively high number and density of electrode array (96 channels). It has been previously suggested that CAR would perform reasonably well if the number and density of the electrodes are high [17]. When using a different decoder, gated recurrent unit (GRU) network [33, 45], we also observed the same trend, which indicates the robust performance of CAR. Another finding was that bipolar referencing yielded the second-highest decoding performance after CAR. Apart from the ability to remove common noise, bipolar referencing could also add more informative features (due to many combinations of electrode pairs) to the decoder. A rather surprising finding was that, although CSD could remove common noise, its decoding performance was lower than the other referencing schemes. This is likely ascribed to the fact that electrodes at the edges (not having four neighbouring electrodes) were excluded, effectively throwing away some information.

We further examined whether the above results were dependent on the choice of the feature. When using gamma-driven LSTM decoder, it turned out that bipolar yielded the highest decoding performance followed by CAR, unipolar, and CSD, respectively. Upon investigation on this phenomena, we found that the gamma features obtained from unipolar and CAR were highly correlated (average CC = 0.78), whereas LMP features were less correlated (average CC = 0.50). This suggests that CAR is not effective when using gamma as the feature. It is important to note that the decoding performance of CAR coupled with LMP was found to be significantly better than bipolar coupled with gamma.

Lastly, we examined the impact of referencing schemes on the interchannel correlation. We observed that bipolar, CSD, and CAR decreased the interchannel correlation substantially with respect to unipolar. However, the decrease in interchannel correlation thought to reflect some degree of common noise does not necessarily translate into better decoding performance. The decoding performance is not only affected by the use of referencing schemes, but also by other factors such as feature type and feature size, among others. For example, as previously described, although CSD had substantially lower interchannel correlation, it also had a smaller number of features, which led to lower decoding performance relative to unipolar.

The impact of referencing schemes on the analysis of LFP signals (e.g. power, phase, coherence, and Granger causality) have been comparatively studied before. Our present study extends these prior studies by comparing the impact of referencing schemes on the decoding performance of LFP-based BMI. To the best of our knowledge, this is the first study that comprehensively investigates the impact of referencing schemes for BMI application using LFPs. Similar studies using EEG signals have reported contradictory findings, that is, CSD (i.e. Laplacian) yielded the best performance [46, 47, 48]. However, it is difficult to make a direct comparison between these studies and ours due to notable differences in several factors such as signal sources (scalp vs intracortical), subjects (human vs monkey), behavioural tasks (movement imagery vs movement execution), signal processing and decoding algorithms, among others.

In summary, our overall results suggest the use of CAR coupled with LMP for LFP-based BMIs. Both CAR and LMP are simple, computationally lightweight, and amenable for on-chip implementation and real-time application [49, 2]. On top of that, LFPs have been demonstrated to exhibit long-term signal stability [1, 4, 5], which is critical for widespread clinical use of a BMI device.

## 5. Conclusion

We have presented a systematic and comprehensive investigation of the impact of the referencing schemes on the decoding performance of LFP-based BMI. Our experimental results revealed that LMP was the most informative LFP feature regardless of the referencing schemes, indicating its richness of movement-related information. Using LMP-driven deep learning algorithms, we further showed that CAR significantly and consistently yielded better decoding performance relative to other referencing schemes. These findings provide empirical justification for the use of CAR coupled with LMP for enhancing the decoding performance of LFP-based BMI. The simplicity and amenability of both CAR and LMP for on-chip implementation along with the long-term stability of LFP signals could potentially pave the way towards low power, fully implantable, and clinically viable BMI.

## Acknowledgment

This work was partially funded by the Engineering and Physical Sciences Research Council (EPSRC) grant EP/M020975/1 and the Indonesia Endowment Fund for Education (LPDP) grant PRJ-123/LPDP/2016. We thank J. E. O’Doherty and P. N. Sabes for making their data publicly available.

